## Supplementary material for "Maximizing CRISPRi efficacy and accessibility with dual-sgRNA libraries and optimal effectors": Supplementary_Note_1.docx

**Supplementary Note 1: Ordering, cloning, and sequencing pooled, programmed dual sgRNA libraries.**

Modified from <https://weissman.wi.mit.edu/resources/Pooled_CRISPR_Library_Cloning.pdf>

**Overview**

This protocol describes the ordering and cloning of pooled, *programmed* dual sgRNA libraries.

As a general strategy, we first order dual-protospacer oligos as an oligonucleotide pool. Oligo pools are PCR-amplified, digested with BstXI/BlpI, gel extracted, and ligated into a similarly digested single sgRNA lentiviral vector. This ligation product is then transformed into bacteria and amplified at scale (>100 bacterial colonies per oligo in the pool) to generate an intermediate library. Next, a constant region-U6 promoter drop-in is BsmBI digested and ligated into the BsmBI-digested intermediate library. The final library is then transformed into bacteria at scale and sequenced.

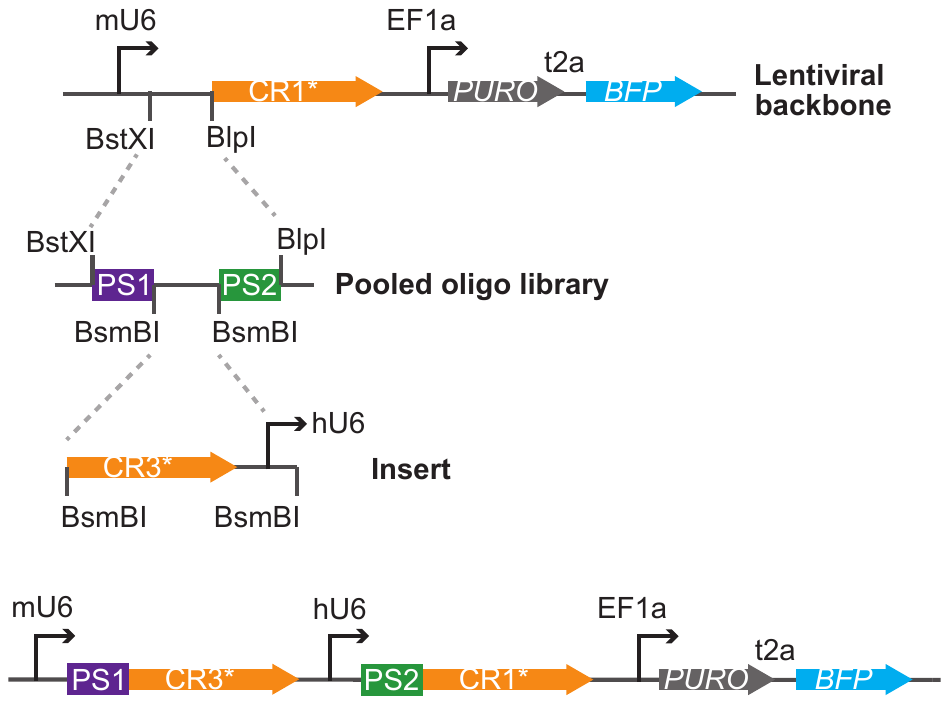

Figure 1. Schematic of library cloning strategy.

**1) Design and ordering oligo libraries from Agilent/Twist/IDT**

For dual sgRNA libraries, we require three elements:

1. A base lentiviral vector. The selection of vector depends on your desired markers and sgRNA constant region. we have vectors with multiple selectable markers. If the goal is to perform 5’ direct capture Perturb-seq or standard growth/sort screens, we clone with vectors that do not contain a capture sequence. The guides without capture sequences have higher expression (see <https://doi.org/10.1101/503367>; Figure 1h). If the goal is to perform 3’ direct capture Perturb-seq, we clone with vectors that contain capture sequences. *Ensure that your vector does not contain BstXI, BlpI, or BsmBI cut sites outside of the guide cassette (these enzymes are used for library cloning).*

| **plasmid** | **U6 promoter** | **constant region** | **capture sequence** | **Pol-II promoter** | **Selectable marker** |
| --- | --- | --- | --- | --- | --- |
| pJR100 | mU6 | modified CR1 | CS1 in stem loop | UCOE-EF1alpha | PURO-BFP |
| pJR101 | mU6 | modified CR1 | CS1 in stem loop | UCOE-EF1alpha | PURO-GFP |
| pJR103 | mU6 | CR1 | none | UCOE-EF1alpha | PURO-BFP |
| pJR104 | mU6 | CR1 | none | UCOE-EF1alpha | PURO-GFP |

2. A dual-guide oligo library.
   a. To order the library, begin by picking sgRNAs from your favorite sgRNA table. For CRISPRi or CRISPRa, we use <https://www.ncbi.nlm.nih.gov/pubmed/27661255>. *Ensure that all sgRNAs have a 5’ “G” for U6 transcription initiation. Ensure that your sgRNAs do not contain BstXI, BlpI, or BsmBI cut sites (these enzymes are used for library cloning).*
   b. Pick PCR adapter sequences for oligo amplification from: <http://weissmanlab.ucsf.edu/CRISPR/library_adapters.xlsx>
   c. Choose whether the constant region CR1, CR2, or CR3 will be used for the first sgRNA. These constant regions are taken from <https://www.ncbi.nlm.nih.gov/pubmed/27984733>. *The choice of constant region will change the sequence of the spacer between the two protospacers.* I generally use a mU6-CR3-hU6-CR1 design, but other designs may be desirable.
   d. Assemble the oligos.

| **design** | **oligo assembly** |
| --- | --- |
| mU6-CR3-hU6-CR1 | 5’- PCR adapter -  CCACCTTGTTG - protospacer A - gtttcagagcgagacgtgcctgcaggatacgtctcagaaacatg - protospacer B - GTTTAAGAGCTAAGCTG - PCR adapter-3’ |
|  | example CR3 oligo: ATTTTGCCCCTGGTTCTTccaccttgttgGGGGCGCTGGAAACCCTTACgtttcagagcgagacgtgcctgcaggatacgtctcagaaacatgGTCCCGCGCCTCCAAAACCTgtttaagagctaagctgCCAGTTCATTTCTTAGGG |
| mU6-CR2-hU6-CR1 | 5’- PCR adapter -  CCACCTTGTTG - protospacer A - gtttgagagcgagacgtgcctgcaggatacgtctcagaaacatg - protospacer B - GTTTAAGAGCTAAGCTG - PCR adapter-3’ |
|  | example CR2 oligo: ATTTTGCCCCTGGTTCTTccaccttgttgGGGGCGCTGGAAACCCTTACgtttgagagcgagacgtgcctgcaggatacgtctcagaaacatgGTCCCGCGCCTCCAAAACCTgtttaagagctaagctgCCAGTTCATTTCTTAGGG |

e. Order the oligo library. We always order the oligos in the forward and reverse complement direction to decrease dropouts during oligo synthesis.

3. A constant region-U6 promoter drop-in. As described above, if the goal is to perform 5’ direct capture Perturb-seq or standard growth/sort screens, we clone with vectors that do not contain a capture sequence. If the goal is to perform 3’ direct capture Perturb-seq, we clone with vectors that contain capture sequences. The drop-in elements are synthesized by IDT as a MiniGene ordered on a pUCIDT ampicillin resistant plasmid.

| **plasmid** | **U6 promoter** | **constant region** | **modifications** |
| --- | --- | --- | --- |
| pJR89 | hU6 | modified CR3 | CS1 in stem loop |
| pJR97 | hU6 | CR2 | none |
| pJR98 | hU6 | CR3 | none |
| pJR98-IBC | hU6 | CR3 | IBC; see addendum |

***For the rest of this protocol, I will describe cloning in pJR104 with a CR3/CR1 oligo pool and pJR98 drop-in.***

### 2) Vector Preparation: pJR104 library vector

a) Digest 5 ug of vector with NEB BstX1 and Blp1 for 2-6h at 37ºC. Set up reaction as follows:

|  | **100 uL Reaction** |
| --- | --- |
| 10x 2.1 Buffer | 10 uL |
| pJR104 vector | (5ug) |
| NEB Blp1 | 4 uL |
| NEB BstX1 | 4 uL |
| Water | Fill to 100 uL |

Depending on the size of your library, you may need to set up multiple reactions. A typical yield ranges from 10%-30%.

b) While digest is incubating, make 0.8% agarose gel and use wide gel combs. Stain gel and excise linearized vector.

c) Gel purify excised vector using an appropriate gel purification kit, (e.g. Machery-Nagel’s NuceloSpin Gel & PCR Cleanup kit (Catalog # 740609), Qiagen, Zymo)

### 3) Insert Preparation: PCR

As a general strategy, the insert PCR has to serve three purposes.

- First, it must amplify sublibraries from the oligo pool such that the sublibrary constitutes the vast majority of the product. I do not recommend performing less than 10 PCR cycles as 2^10=1024.
- Second, it must produce sufficient material for the digestion and ligation steps. 1 ug is ideal.
- Third, it should avoid introducing excessive bias and heteroduplex/bubble products caused by overamplification. We check for heteroduplexes by looking for higher molecular weight species on the bioanalyzer or gel (if you want to prove that a band is a heteroduplex, perform a reconditioning PCR: ncbi.nlm.nih.gov/pmc/articles/PMC113844/). As a general rule, heteroduplexes indicate that the library is overamplified and I tend to redo the PCR with less cycles to minimize bias.

Depending on the number of sublibraries and available template material, you may need to adjust the PCR conditions. I often perform a test PCR to find a condition which satisfies the above criteria before using excessive template material.

a) Dilute oligo library in Buffer EB to approximately 2 ng/uL (~0.04 pmol/uL assuming ssDNA dual-guide oligos are 148 bp).

- Twist ships libraries with yields depending on the number of elements.
- IDT ships 10 pmol per library. As 10 pmol = 457.5 ng, then add 228.75 uL Buffer EB to dilute to 2 ng/uL.

b) Set 4 X 100 uL reactions and an additional “NO template” negative control. While 4 X 100 uL reactions is excessive you will have plenty of back-up insert.

| **reagent** | **100 uL test** | **400 uL final** |
| --- | --- | --- |
| Water | 61 | 244 |
| 5x Phusion HF Buffer | 20 | 80 |
| 100% DMSO | 3 | 12 |
| dNTPs (10mM ea) | 2 | 8 |
| Forward Primer (10 uM) | 5 | 20 |
| Reverse Primer (10 uM) | 5 | 20 |
| **TEMPLATE oligo pool (2 ng/uL)** | 3 | 12 |
| HF Phusion | 1 | 4 |
| Total | 100 | 400 |

| **# Cycles** | **Temp** | **Time** |
| --- | --- | --- |
| 1 | 98C | 30sec |
| 11X | 98C | 15sec |
|  | 56C | 15sec |
|  | 72C | 15sec |
| 1 | 72C | 10min |
| hold | 4C | inf |

c) Combine reactions in a 5 mL tube (or similar). To verify correct amplification, run 5 uL of combined reactions on a 10% acrylamide gel at 120V for ~20-30 minutes. Alternatively, run a BioAnalyzer High Sensitivity DNA chip.

d) While gel/BioAnalyzer is running purify the remaining ~395 uL of each PCR using Qiagen’s MinElute Reaction Cleanup Kit (or similar) (Catalog #28204), using 1 column per pooled sample:

- Add 4X sample volume of Qiagen Binding Buffer PB to combined PCR.
- Add 1/100th volume 3M NaOAc.
  1. Combined PCR= ~350 uL, Add 1400 uL PB and 17.5 uL NaOAc
- Once PCR is bound to column, Wash 2X w/ 750 uL of PE
- Perform one dry spin at max speed for 2 minutes
- Elute in 20 uL of EB (warm media to 70C and wait 5 minutes before eluting)

*NOTE: The MinElute Kit does not come with Buffer PB for our high-volume libraries, so additional Buffer PB must be purchased separately (Catalog #19066).

Your recovery should be between ~0.3 ug to 1 ug (4,100 uL rxns). If your recovery is lower than 0.3 ug, you will need to optimize your PCRs before proceeding with the digest**. I generally quantify by Qubit.**

e) At this point the gel/BioAnalyzer should be finished. **The expected insert size is 148 bp.** Once you have verified your Insert PCR is correct and you do not observe excessive heteroduplex/bubble PCR product, you can proceed and digest your insert.

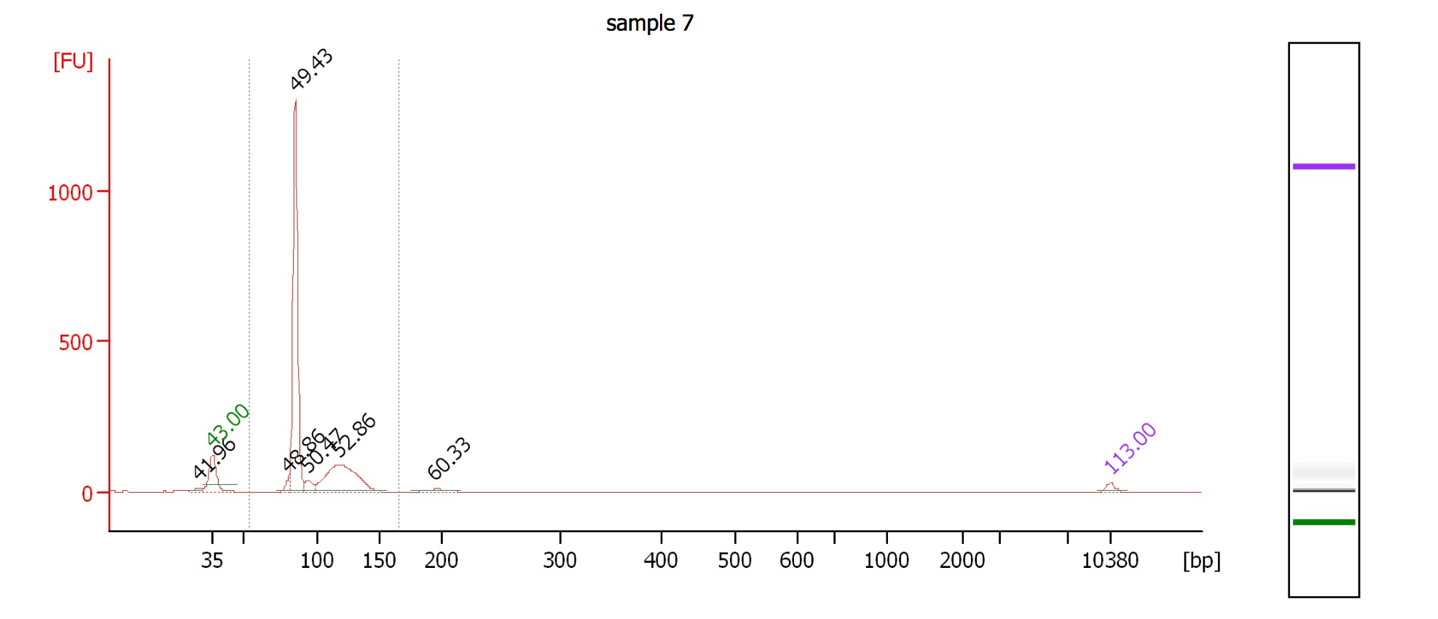

This figure shows an example of a overamplified single-sgRNA library. The desired peak is observed at 84 bp. The overamplification leads to a heteroduplex peak at ~120 bp. The gel image on the right shows how this would appear on a gel. This PCR could be repeated with less cycles to minimize PCR bias.

**4) Insert Digest:**

a) Digest 1 ug of purified insert**.** Incubate digest at 37ºC for 2-6 hours. Following digest insert will be ~97bp (148=26+97+25).

| Component | Volume |
| --- | --- |
| Insert (~1 ug) | 20 |
| BstXI | 2 |
| BlpI | 2 |
| 10X 2.1 Buffer | 3 |
| H20 | 2 |
| Total | 30 |

b) Gel purification

- To purify, run the digest on a 10% acrylamide gel. Load 15 uL of digest into each of 2 adjacent wells, leaving an empty spacer well between different samples. Run the gel at ~120V for ~30-40 minutes, or until the yellow dye has traveled at least 75% of the total lane length
- While gel is running prepare tubes and tips for rapid extraction:
- When the gel has completed running, stain with EtBr or SYBRsafe and excise insert using UV or blue light transilliminator. Change blades between samples to avoid cross contamination. Place excised gel piece inside 0.5mL nonstick tube prepared above, and proceed with Rapid Extraction.

c) Rapid Extraction

- Pierce a 0.5mL nonstick tube (Ambion, Catalog #AM12350) with an 18.5 gauge needle and place inside a 1.5 mL nonstick tube
- Excise gel piece and place it inside of 0.5ml tube with a hole.
- Spin tubes at 20,000 x g for 3 min. This will crush the gel as it is forced through the hole. Check for gel pieces in the 0.5 mL tube. Residual pieces should be transferred to large tube. If you cannot tap the residual pieces use a pipette tip.
- Add 200 uL of water to gel pieces, and incubate for 30min~1 hour at 70ºC.
- Vortex gel slurry for 30s and use cut p1000 tips to transfer gel mixture to Costar Spin-X columns.
- Spin tubes for 3 min at 20,000 x g to recover the elution mixture free of gel debris.
- Transfer eluate to a new 1.5 mL nonstick tube
- Ethanol Precipitation
  - 200 uL of eluate (insert is here)
  - 2uL of glycoblue (ThermoFisher #AM9515) *this will help visualize your pellet. Vortex thoroughly before adding to each sample.
  - 20 uL 3M NaOAc or 3M NaCl **Mix Well (invert 10-15X)**.
  - Add 0.6 mL 100% EtOH, mix well
  - Incubate at -20ºC, for 2hr to overnight.
  - Pellet: Spin 45 min at 20,000 x g at 4ºC , remove supernatant
  - Wash pellet 2X with ice cold 80% EtOH
  - Air Dry
  - Resuspend pellet in 25 uL water or Qiagen EB

d) If you nanodrop your sample you will observe a contamination peak at 230. This is from the glycoblue. I generally observe a nanodrop concentration of ~20-25 ng/uL with glycoblue, and without glycoblue ~10-15 ng/uL. From Qubit, a fluorometer, (DNA high sensitivity assay) I observe ~0.5-2 ng/uL. As the Qubit is more sensitive I use these concentrations. Proceed to ligation.

### 5) Intermediate Ligation

As a first pass, you may want to set up 3 conditions (vector:insert), 1:2, 1:1, 2:1. Adding too much insert can cause sgRNA concatemers, which are observed at 1:3. The molar ratios of 1:2 worked well for me (500 ng vector:10.6 ng insert).

|  | 20 uL ligation |
| --- | --- |
| Vector | (500 ng) |
| Insert | (10.6 ng) |
| 10x Ligase buffer | 2 uL |
| T4 ligase | 1 uL |
| Water | Fill to 20 uL |

**Include a negative control with no insert to check for background from vector.**

**Incubate at 16C for 16 hours.**

I generally use the NEB T4 ligase with 5X activity units per volume (catalog #M0202T/M0202M). However, this is not strictly necessary. A 500 ng ligation is normally sufficient to yield up to 10 million colonies, but this can depend on vector/insert purity and proper ratio.

a) **Ligation Reaction Clean-Up: EtOH Precipitation**

*Removing the excess salt in the ligation reaction is crucial for electroporation.*

**EtOH Precipitation** (alternatively, perform a 2X SPRI eluting in water)

- Bring 20 uL ligation reaction to 200 uL with water.
- Add 2uL of glycoblue (this will help you visualize your pellet)
- ● Add 1/10 reaction volume of 3M NaOAc, 20 uL. **Mix Well.**
- Add 3X reaction volume of 100% EtOH, 600 uL. Invert 10X to mix.
- Place at -20ºC, 2h to overnight.
- Pellet: Spin 45 min at 20,000 x g at 4ºC
- Remove supernatant and wash pellet 2X with ice cold 80% EtOH
- Air Dry
- Resuspend pellet in 20 uL of water
- From a 20 uL ligation, we ideally observe ~15 ng/uL (from Qubit). Unfortunately, the EtOH precipitation yields can be variable

#### 6) Test Transformation

I perform a test transformation followed by colony PCR and Sanger sequencing to verify correct inserts and successful ligations with low background.

a) Primary transformation with high efficiency chemically competent cells (ex, Stellar cells (Clontech, # 636736, 108),

- Add 0.5uL ligation to 20 uL Stellar cells, and gently tap to mix.
- Incubate on ice for 30 minutes
- Heat-shock 45s @ 42ºC
- Incubate on ice for 2 min
- Add 300 uL pre-warmed SOC and incubate for 1 hour at 37ºC.
- Plate 200 uL on pre-warmed plates (save the remaining recovery at RT or 4C). Additionally, add 100 uL to a 3 mL liquid culture of LB+carb for plasmid prep.
- Incubate O/N at 37ºC

b) Plasmid prep and sanger sequence:

If your negative control shows <5% background colonies, it is safe to proceed. Purify plasmid by mini-prep from liquid cultures. Sanger sequence the plasmid library to ensure the expected insert is observed (with mixed peaks in the two protospacer positions).

c) Colony PCR:

- Expected length of dual-guide insert product in CR1/no capture sequence vector: 285 bp
- Expected length of single-guide insert product OR reanealled vector (no capture sequence): 221 bp
- Expected length of dual-guide insert product in CR1-CS1 vector: 311 bp
- Expected length of single-guide insert product OR reanealled vector (CR1-CS1): 247 bp
- This primer pair will not bind to the CROPSEQ (pJR107 - dJR004 and dJR009).

| oJR251 | cagcacaaaaggaaactcacc |
| --- | --- |
| oJR252 | cgcctaatggatcctagtact |

| **Component** | **50 µl Reaction** |
| --- | --- |
| Q5 High-Fidelity 2X Master Mix | 25 |
| 10 µM Forward Primer | 2.5 |
| 10 µM Reverse Primer | 2.5 |
| Template DNA (1:100) | 1 |
| Nuclease-Free Water | 20 |
| Total | 50 |

| Initial denaturation | 98˚ | 30 sec |
| --- | --- | --- |
| 10X | 98˚ | 10 sec |
|  | 64˚ | 30 sec |
|  | 72˚ | 20 sec |
| final extension | 72˚ | 2 min |
|  | 4˚ |  |

Dilute 1:5 and then bioanalyze to check for desired length without concatamers.

#### 7) Large-Scale Transformation

- For the large-scale transformation, it is necessary to maintain at least 30X coverage of the library. Thus, for a 1000 element library, we obtain a minimum of 30000 colonies to maintain diversity. To minimize variation in the library for smaller libraries (e.g. Perturb-seq libraries), I typically clone at 100X coverage. For libraries containing IBCs, I generally only use 10X coverage per sgRNA-IBC combination.
- I typically plate 1/1000 of the total recovery media on a 10 cm LB+carb plate in order to count colonies and transfer the remaining recovery media directly into liquid culture (1 ml recovery media into ~100 ml LB+carb).
- We want to use at least 50 uL of MegaX cells for every 10,000 library elements. I typically combine 10 uL of EtOH-precipitated ligation product with 50 uL of MegaX cells. Each electroporation cuvette holds a maximum of 90 uL. Thaw Mega-X cells on ice and pre-chill cuvettes on ice.

a) Transformation protocol:

- In a 1.5 mL eppendorf tube, mix appropriate volume of Mega-X and ligation product

(~100 ng of ligation per 20 uL of MegaX). Tap gently 3X to mix.

- Incubate on ice for 30 min
- Transfer the mix to prechilled cuvette using long gel-loading tips. Be sure there are no bubbles in your MegaX/dna mix
- Electroporate at 2.0 kV, 200 ohms, 25 uF in 0.1 cm cuvette (Gene Pulser Xcell, Bio Rad) You should observe a TC between 4.4ms and 4.8ms (this depends upon volume; values above 5ms are likely not successful electroporations). You may need to use less ligation per cuvette reaction, and/or double check for bubbles.
- Following electroporation, rinse each cuvette twice with 600 uL of recovery media or S.O.C and transfer to prepared cell culture tube. Recover at 37C at ~250RPM for 2hrs.
- Plate 1/1000 of recovery media for colony counting. Combine remaining recovery media with LB+carb (1 mL: 100 mL) for overnight culture.
- Incubate at 37ºC for 18h and make sure that colonies are grown well on all plates.
- The next day, spindown 50 mL of overnight culture for midi prep.

***This intermediate library now must go through another digestion and ligation in order to obtain a functional dual-sgRNA library. The length of the digested intermediate library should be ~9kb). The length of the digested pJR89/pJR97/pJR98 inserts is ~400 bp.***

### 8) Vector Preparation #2: intermediate library digestion

a) Digest 5 ug of intermediate library with NEB BsmBI for 2-6h at **55ºC**. Set up reaction as follows:

|  | **100 uL Reaction** |
| --- | --- |
| 10x 3.1 Buffer | 10 uL |
| intermediate library | (5ug) |
| NEB BsmBI | 5 uL |
| Water | Fill to 100 uL |

Depending on the size of your library, you may need to set up multiple reactions. A typical yield ranges from 10%-30%.

b) While digest is incubating, make 0.8% agarose gel and use wide gel combs. Stain gel and excise linearized vector (9kb).

c) Gel purify excised vector using an appropriate gel purification kit, (e.g. Machery-Nagel’s NuceloSpin Gel & PCR Cleanup kit (Catalog # 740609), Qiagen, Zymo).

### 9) Vector Preparation #2: intermediate insert digestion

a) Digest 5 ug of intermediate library with NEB BsmBI for 2-6h at **55ºC.** Set up reaction as follows:

|  | **100 uL Reaction** |
| --- | --- |
| 10x 3.1 Buffer | 10 uL |
| pJR98 | (5ug) |
| NEB BsmBI | 5 uL |
| Water | Fill to 100 uL |

Depending on the size of your library, you may need to set up multiple reactions. A typical yield ranges from 10%-30%.

b) While digest is incubating, make 0.8% agarose gel and use wide gel combs. Stain gel and excise linearized vector **(~400 bp – do not let this run off the gel).**

c) Gel purify excised vector using an appropriate gel purification kit, (e.g. Machery-Nagel’s NuceloSpin Gel & PCR Cleanup kit (Catalog # 740609), Qiagen, Zymo).

### 10) Final Ligation

As a first pass, you may want to set up 3 conditions (vector:insert), 1:2, 1:1, 2:1. The molar ratios of 1:1 worked well for me (500 ng vector: 22 ng insert). If you find that the vector control background colony rate is high, then phosphatase treat the vector prior to ligation.

|  | 20 uL ligation |
| --- | --- |
| Vector | (500 ng) |
| Insert | (22 ng) |
| 10x Ligase buffer | 2 uL |
| T4 ligase | 1 uL |
| Water | Fill to 20 uL |

**Include a negative control with no insert to check for background from vector.**

**Incubate at 16C for 16 hours.**

I generally use the NEB T4 ligase with 5X activity units per volume (catalog #M0202T/M0202M). However, this is not strictly necessary. A 500 ng ligation is normally sufficient to yield up to 10 million colonies, but this can depend on vector/insert purity and proper ratio.

a) **Ligation Reaction Clean-Up: EtOH Precipitation**

*Removing the excess salt in the ligation reaction is crucial for electroporation.*

**EtOH Precipitation** (alternatively, perform a 2X SPRI eluting in water)

- Bring 20 uL ligation reaction to 200 uL with water.
- Add 2uL of glycoblue (this will help you visualize your pellet)
- ● Add 1/10 reaction volume of 3M NaOAc, 20 uL. **Mix Well.**
- Add 3X reaction volume of 100% EtOH, 600 uL. Invert 10X to mix.
- Place at -20ºC, 2h to overnight.
- Pellet: Spin 45 min at 20,000 x g at 4ºC
- Remove supernatant and wash pellet 2X with ice cold 80% EtOH
- Air Dry
- Resuspend pellet in 20 uL of water
- From a 20 uL ligation, we ideally observe ~15 ng/uL (from Qubit). Unfortunately, the EtOH precipitation yields can be variable.

#### 11) Large-Scale Transformation

- For the large-scale transformation, it is necessary to maintain at least 30X coverage of the library. Thus, for a 1000 element library, we obtain a minimum of 30000 colonies to maintain diversity. To minimize variation in the library for smaller libraries (e.g. Perturb-seq libraries), I typically clone at 100X coverage.
- I typically plate 1/1000 of the total recovery media on a 10 cm LB+carb plate in order to count colonies and transfer the remaining recovery media directly into liquid culture (1 ml recovery media into ~100 ml LB+carb).
- We want to use at least 50 uL of MegaX cells for every 10,000 library elements. I typically combine 10 uL of EtOH-precipitated ligation product with 50 uL of MegaX cells. Each electroporation cuvette holds a maximum of 90 uL. Thaw Mega-X cells on ice and pre-chill cuvettes on ice.

a) Transformation protocol:

- In a 1.5 mL eppendorf tube, mix appropriate volume of Mega-X and ligation product

(~100 ng of ligation per 20 uL of MegaX). Tap gently 3X to mix.

- Incubate on ice for 30 min
- Transfer the mix to prechilled cuvette using long gel-loading tips. Be sure there are no bubbles in your MegaX/dna mix
- Electroporate at 2.0 kV, 200 ohms, 25 uF in 0.1 cm cuvette (Gene Pulser Xcell, Bio Rad) You should observe a TC between 4.4ms and 4.8ms (this depends upon volume; values above 5ms are likely not successful electroporations). You may need to use less ligation per cuvette reaction, and/or double check for bubbles.
- Following electroporation, rinse each cuvette twice with 600 uL of recovery media or S.O.C and transfer to prepared cell culture tube. Recover at 37C at ~250RPM for 2hrs.
- Plate 1/1000 of recovery media for colony counting. Combine remaining recovery media with LB+carb (1 mL: 100 mL) for overnight culture.
- Incubate at 37ºC for 18h and make sure that colonies are grown well on all plates.

The next day, spindown 50 mL of overnight culture for midi prep.

#### 12) To prepare samples for Illumina sequencing, see the attached “Illumina Sequencing Sample Prep” guide.

Illumina sequencing of the resulting plasmids is necessary to ensure the resulting plasmid libraries from this protocol are correct, have a large fraction of sgRNAs that match perfectly to the expected library, and the relative fractions of each sgRNA are tightly distributed. You only need ~100 sequencing reads per sgRNA in the library to be able to validate the library.

### Addendum: Arrayed cloning of IBCs into pJR98

### In order to maintain extremely even coverage of the IBC library and ensure that there were no sequence errors in the library, I cloned the library in an arrayed format. For IBC sequences, see: IBC_sequenes_filtered.csv.

### 1) Digest pJR98 with AscI in Buffer Cutsmart

### 2) Gel extract (only one 3.2 kb fragment should be observed on the gel)

### 3) Gibson Assembly. Incubate the assembly reaction for 1 hour at 50°C. Then transform, grow overnight, miniprep, and Sanger sequence with M13/pUC Forward primer (CCCAGTCACGACGTTGTAAAACG).

|  | ssDNA insert (1:100 dilution of 100 uM ssDNA) | vector only control |
| --- | --- | --- |
| 10 ng digested vector (10 ng/uL) | 1 | 1 |
| insert | 1 | 0 |
| NEBuilder HiFI DNA Assembly Master Mix | 10 | 10 |
| ddH20 | 8 | 9 |
| total | 20 | 20 |
