## Supplementary material for "Maximizing CRISPRi efficacy and accessibility with dual-sgRNA libraries and optimal effectors": Supplementary_Note_2.docx

### Supplementary Note 2: Illumina Sequencing Sample Preparation for CRISPRi/a dual-guide IBC libraries

Adapted from <https://weissman.wi.mit.edu/resources/IlluminaSequencingSamplePrep.pdf>

**Overview**

This protocol describes the three steps required for generating sequencing samples from cells harvested from screens conducted with the CRISPRi/a-v3 libraries:

1. Isolation of genomic DNA from fresh or frozen cell pellets
2. PCR of sgRNA-containing fragments to amplify and append Illumina sequencing adapters
3. PCR clean-up and submitting for sequencing

Note: Previous versions of this protocol included an additional gel extraction step between steps 1 and 2. With newer PCR products capable of handling large amounts of genomic DNA inputs, this step can now be omitted, at the cost of setting up ~100 PCRs per sample. A version of the protocol including gel extractions is available at: <https://weissmanlab.ucsf.edu/CRISPR/IlluminaSequencingSamplePrep_old.pdf>

### Reagents, kits, and resources

- Macherey Nagel Blood
  - Mini, L, or XL depending on number of cells; if XL, 1 column per 100 million cells
- NEBNext Ultra II Q5 MasterMix (NEB)
  - expect to use ~1 kit of 250 reactions per 100 million cells, depending on yields from preceding steps
- Custom PCR and sequencing primers (sequences in protocol)
- SPRIselect beads (BeckmanCoulter; expensive but pre-made) or Seramag speed beads (Thermo scientific; much cheaper but requires some preparation steps)
- Magnetic stand
- Access to Bioanalyzer (Agilent) with DNA High Sensitivity chips and reagents
- Access to an Illumina sequencing facility

### A word of warning on the sample prep

The sgRNA content of the sample up to the PCR step is at extremely low abundance compared to plasmid or post-PCR DNA! This makes the samples very susceptible to contamination which can be further amplified in the PCR. We do all of the genomic extractions, digests, PCR set-up in a separate room to avoid contamination. You don’t have to be that extreme, but do use a separate bench, set of pipettes, etc. from where you clone sgRNA plasmids or do the post-PCR clean-up step.

### Step 1: gDNA isolation

*We use Nucleospin Blood kits from Macherey Nagel sold in mini, midi(L), and maxi(XL) form.*

- mini: 5 x 10^6,^, midi: 2 x 10^7^; maxi: 1 x 10^8^

1a. For MN NucleoSpin Blood XL prep preheat water bath to 56C, and 70C 2b. Frozen Cells:

1. Thaw cells: RT or 37C
2. Resuspend cells in 25 mL of PBS or transfer cells to 50 mL falcon tube with 25 mL PBS.
3. Pellet cells. Spin @ 400 x g, 5 min

*or*

2c. Freshly Harvesting Cells

1. Spin @ 400 x g, 5 min, and remove supernatant.
2. Add 25 mL of PBS to 50 mL falcon tube. Transfer cells to 50 mL falcon tube.
3. Pellet cells. Spin @ 400 x g, 5 min
3. **Macherey Nagel (MN) NucleoSpin Blood** (follow NucleoSpin Blood protocol and guide below; detailed protocol provided as ):

| **Number of cells*:** | **< 100M** | **< 200M** | **< 300 M** |
| --- | --- | --- | --- |
| Resuspend in PBS | 10 ml | 20 ml | 30 ml |
| Add ProK | 500 ul | 1,000 ul | 1,500 ul |
| Add BQ1 | 10 ml | 20 ml | 30 ml |
| Add 100% EtOH | 10 ml | 20 ml | 30 ml |
| Split among how many columns: | 1 | 2 | 3 |

- Round up for highest recovery. 150M use 2 columns; 250M, use 3 columns... Notes:
  - After incubating cells at 56C, remember to cool lysed cells to RT before adding 100% EtOH. You can place tubes on ice for 5-10min but ideally leave them to cool at room temperature for ~60 min.
  - MN recommends spinning @ 4000x g; if your centrifuge cannot reach this, double spin times to maximize recovery
  - Prior to eluting, preheat EB at 70C. Add 0.8 mL EB per column, incubate at RT 5 min, spin 4,000 rpm 4 min. Repeat. Recovery per column is roughly 1.2-1.6 mL.
  - You should recover ~ 1-1.5 mg of gDNA per column

### Step 2: PCR for sgRNA amplicon enrichment and Illumina adapter ligation

The PCR is designed to enrich (amplify) dual-sgRNA constructs. During the enrichment, Illumina P5 and P7 adapters are added to the 3’ and 5’ end of the sgRNA cassette, and the index (barcode for de-multiplexing) is also added to one end. Below is a cartoon depicting the reaction, followed by sequences of our primers. We order the primers from IDT with standard desalting.

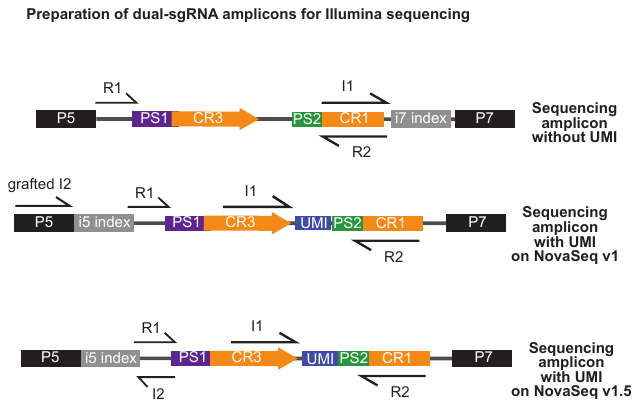

### Note: This protocol is optimized for sequencing on the Illumina MiSeq or NovaSeq 6000. If you are using other sequencers, consult your core facility or test the requirements for sequence diversity. The amplicon for sequencing on MiSeq/NovaSeq v1 is the same as the amplicon for sequencing on NovaSeq v1.5. However, there are different custom sequencing primers for these platforms, depending on the use of a grafted P5. See below.

If your library has sufficient base diversity (for instance, >100 sgRNAs), then it is not necessary to spike phi-X into the sequencing reaction. However, if you are unsure, then spike-in 5-10% phi-X and ensure that the custom sequencing primers are added to the Illumina sequencing primer wells (as opposed to custom sequencing primer wells). If you extend the length of the reads into the constant sgRNA backbone, it is necessary to spike-in phi-X to ensure base diversity.

| **primer** | **Sequence (5’->3’)** |
| --- | --- |
| oJR437 (read 1 sequencing primer) | CGCGTATCCCTTGGAGAACCACCTTGTTGG |
| oJR438 (index read 1 sequencing primer for IBC) | TGCTCGAATCTACACTCAGCTATGGCGCTG |
| oJR439 (read 2 sequencing primer) | GCGGCCGGCTGTTTCCAGCTTAGCTCTTAAAC |
| oJR440 (index read 2 sequencing primer for index; only used for NovaSeq v1.5) | CCAACAAGGTGGTTCTCCAAGGGATACGCG |
| oJR441 (constant reverse CR1 primer) | CAAGCAGAAGACGGCATACGAGATGCGGCCGGCTGTTTCCAGCTTAGCTCTTAAA |
| oJR442 (example i5 index primer, N701 index) | AATGATACGGCGACCACCGAGATCTACACtcgccttaCGCGTATCCCTTGGAGAACCACC |

**Table of indexed forward PCR primers.**

The index that is appended to sgRNA samples allows each sample to be uniquely identified, even when pooled into a single sequencing lane. All indexes below have a Hamming distance of at least 2. Consult with your sequencing facility to find out how many reads to expect per lane, and plan for 1000 reads per sgRNA per sample. For example, a typical genome-scale 20k library screen should have at least 20M reads per sample (e.g. T0, treated, untreated), and each should be PCR amplified such that they have a unique index. Before proceeding plan out how you want to pool/run your samples on the flow cell. If you are sharing a lane with another lab member ask them which indices they are using.

| **primer** | **Sequence (5’->3’, index in lowercase)** | **i5 index for sample sheet** |
| --- | --- | --- |
| oJR442 (i5 N701) | AATGATACGGCGACCACCGAGATCTACACtcgccttaCGCGTATCCCTTGGAGAACCACC | TAAGGCGA |
| oJR443 (i5 N702) | AATGATACGGCGACCACCGAGATCTACACctagtacgCGCGTATCCCTTGGAGAACCACC | CGTACTAG |
| oJR444 (i5 N703) | AATGATACGGCGACCACCGAGATCTACACttctgcctCGCGTATCCCTTGGAGAACCACC | AGGCAGAA |
| oJR445 (i5 N704) | AATGATACGGCGACCACCGAGATCTACACgctcaggaCGCGTATCCCTTGGAGAACCACC | TCCTGAGC |
| oJR446 (i5 N705) | AATGATACGGCGACCACCGAGATCTACACaggagtccCGCGTATCCCTTGGAGAACCACC | GGACTCCT |
| oJR447 (i5 N706) | AATGATACGGCGACCACCGAGATCTACACcatgcctaCGCGTATCCCTTGGAGAACCACC | TAGGCATG |
| oJR448 (i5 N707) | AATGATACGGCGACCACCGAGATCTACACgtagagagCGCGTATCCCTTGGAGAACCACC | CTCTCTAC |
| oJR449 (i5 N708) | AATGATACGGCGACCACCGAGATCTACACcctctctgCGCGTATCCCTTGGAGAACCACC | CAGAGAGG |
| oJR450 (i5 N709) | AATGATACGGCGACCACCGAGATCTACACagcgtagcCGCGTATCCCTTGGAGAACCACC | GCTACGCT |
| oJR451 (i5 N710) | AATGATACGGCGACCACCGAGATCTACACcagcctcgCGCGTATCCCTTGGAGAACCACC | CGAGGCTG |
| oJR452 (i5 N711) | AATGATACGGCGACCACCGAGATCTACACtgcctcttCGCGTATCCCTTGGAGAACCACC | AAGAGGCA |
| oJR453 (i5 N712) | AATGATACGGCGACCACCGAGATCTACACtcctctacCGCGTATCCCTTGGAGAACCACC | GTAGAGGA |

### gDNA PCR

Expect to set up between 15 and 50 PCRs per 25M cell sample, depending on the yield of the genomic DNA extraction. We do not recommend exceeding an input of 10 ug gDNA per 100 uL reaction, as excess template DNA can lead to PCR-based uncoupling of guides (Hegde *et al*. 2018, PMID 29799876). Because this is a new protocol, I recommend doing a test single reaction PCR to check for efficient amplification before using all of your genomic DNA.

### PCR parameters

| Reagent | uL per 100 uL reaction |
| --- | --- |
| oJR441 reverse primer (100 uM) | 1 |
| Indexed forward primer (100 uM) | 1 |
| NEBNext Ultra II Q5 MasterMix | 50 |
| gDNA | 10 ug |
| Water | bring to final volume of 100 uL |

**Protocol**

1) Measure gDNA concentration and determine # of reactions to set up (10 ug per PCR)
2) Transfer gDNA into a 5 mL Eppendorf or a 15 mL Falcon tube (should have ~1.2-1.6 mL from one gDNA extraction column).
3) Add corresponding common and indexing primers (1 uL per reaction).
4) Bring to half the final volume with water.
5) Add corresponding volume of 2x PCR MasterMix.
6) Transfer into a clean, segmented multichannel reservoir and then transfer into a 96-well PCR plate using a multichannel pipette.
7) Run PCR using cycling conditions below.

### PCR conditions

- 98 ºC, 30 s
- 22-24 cycles (see note below)
  - 98 ºC, 10 s
  - 63 ºC, 75 s
- 72 ºC, 5 min
- 4 ºC hold

Note on number of cycles: empirically, 22 cycles produce sufficient amount of product. To be absolutely certain that enough PCR product will be obtained, cycle number can be increased to 23 or 24, with the caveat that each additional cycle may induce non-linearities or PCR uncoupling.

After the PCR is completed, pool all your reactions for a given sample in a 5mL or 15 mL tube. Mix reaction well. All of the sgRNAs in the pool should be represented at much greater abundance than, so you may proceed with only a fraction of the sample for PCR purification. Remember that this material represents a significant contaminant to pre-PCR samples, so be sure to keep them separate.

### Step 3a: PCR purification

*There are several possible ways to remove the gDNA and primer background from the PCR. The recommended method is listed below, but two other options are provided in Appendix 2.*

### Note: with this protocol there will be a large amount of primers in the PCR. We generally recommend performing two rounds of purification to remove excess primer contamination before sequencing.

**SPRI bead selection. Proceed with 100-300 uL of pooled PCR**

For dual-guide libraries, the expected amplicon is 562 bp for guides without Perturb-seq capture sequences. For guides with capture sequences, the expected amplicon is 600 bp. In both cases, we perform a 0.5-0.65X SPRI to clean up the PCR and bioanalyze a 1:5 dilution.

You will perform a double SPRI purification using SPRIselect Reagent (Beckman and Coulter) to select for your 562 bp sample. Double SPRI allows you to exclude anything above and below your target size. You will need: low retention/nonstick tubes, magnetic stand, 80% EtOH (make fresh), and Elution Buffer.

Some references if you are curious about how SPRI beads work:

Fisher S, *et al.*: A scalable, fully automated process for construction of sequence-ready human exome targeted capture libraries. *Genome Biol* 2011

DeAngelis MM, Wang DG, Hawkins TL: Solid-phase reversible immobilization for the isolation of PCR products. *Nucleic Acids Res* 1995

Lis JT. Size Fractionation of double stranded DNA by precipitation with polyethylene glycol. *Nucleic Acids Research* (1975)

Amounts listed below are for 300uL of PCR reaction, but you can adjust volumes according to the input.

1) Add 0.5X SPRI beads.
   - 300 uL * 0.5= 150uL SPRIselect. At this ratio, fragments >650 will bind to the beads
   - Mix well. You want a homogenous mixture. If you splash any of the mixture onto the sides of your tubes or cap spin down briefly.
   - Incubate 10 min at RT
   - Place tubes on magnetic stand for 5 min or until supernatant clears
   - Transfer the supernatant to a new tube. **KEEP supernatant**. Your sample is here.
2) Add 0.65X SPRI beads
   - 300 uL * 0.65 = 195 uL SPRIselect. At this ratio, fragments >150 will bind to the beads.
     - **Note this is based on the original PCR input volume, not the volume of the supernatant obtained from the first SPRI selection. As the supernatant already contains 150 uL of SPRIselect reagent (without beads), you only add 45 uL additional SPRIselect reagent (now with beads) to bring to 0.65X.**
   - Mix well and Spin tubes briefly.
   - Incubate 10 min at RT
   - Place tubes on magnetic stand for 5 min or until supernatant clears
   - Remove supernatant. **KEEP beads**. Your sample is bound to the beads.
   - Rinse beads with 1mL (make sure all your beads are covered) fresh 80% EtOH. Wait 2 min. Remove EtOH. Repeat for a total of 2 washes.
   - Air Dry 2-5 min. The beads will look glossy after your remove the EtOH. The texture will turn from glossy (wet) to matte (dry).
   - Once dry, elute in 30 uL of EB.

### STEP 3b: Sequencing library QC and submission for sequencing

Once Your PCR Product is Purified:

1) Measure PCR clean-up yield by Nanodrop and/or Qubit, and dilute a small portion of your sample to 400 pg/uL based on this measurement.
2) Run samples on a Bioanalyzer (Agilent) with High Sensitivity DNA chip. The enriched PCR product is at 562 bp, but the bioanalyzer peak will run in around 555bp-580bp. You can pool multiple samples together prior to analyzing, or analyze each sample in a separate channel to obtain a more accurate concentration of the sequencing sample prior to pooling.

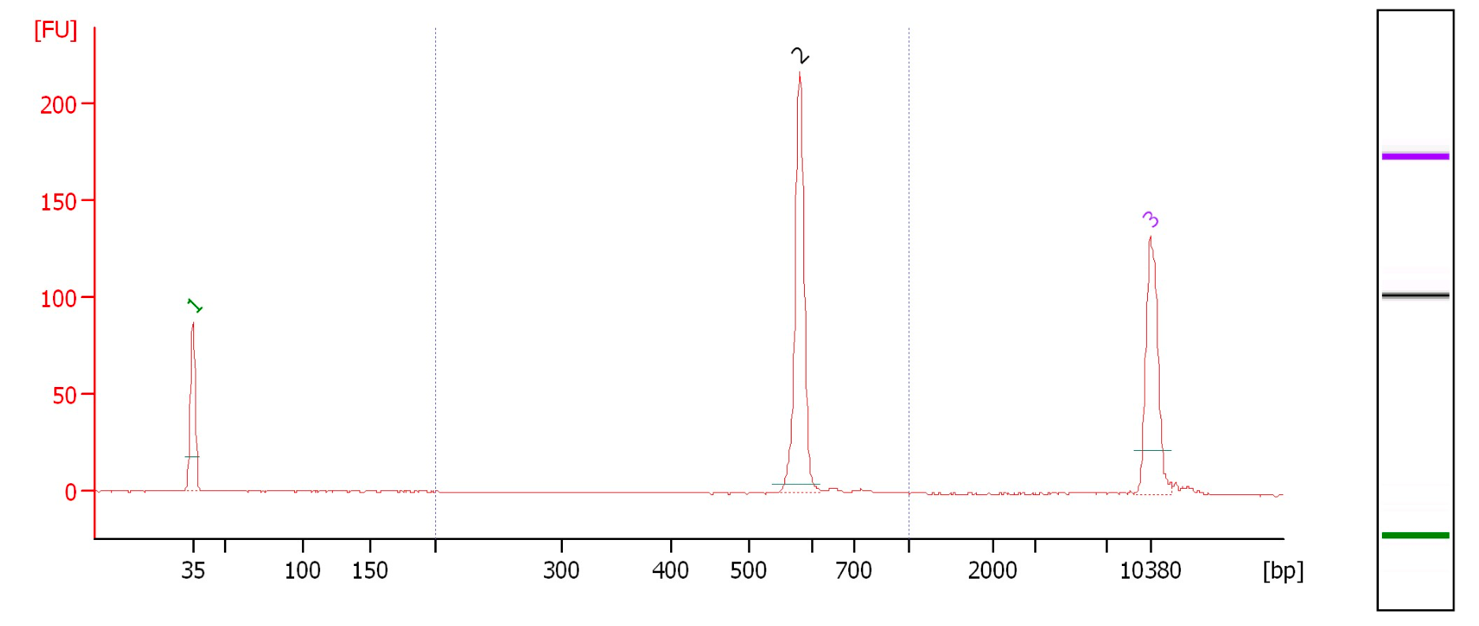

Your sequencing facility may request these traces, and also request or perform additional QC in the form of qPCR.

3) Submit samples for sequencing, along with the custom sequencing primer listed in the PCR section. The exact input volume and molar concentration will depend on the preferences of your sequencing facility. Our facility requests 15uL of 3-10nM (1-3ng/uL) pooled sample, with an additional 5uL for each lane.

The sequencing parameters are:

### Run type: Paired-end 50 with R1: 19, I1: 8, R2: 19, I2: 8.

- Sequencing primers: Read 1 custom primer oJR437; Index Read 1 custom primer oJR438; Read 2 custom primer oJR439.
- If you are sequencing on a NovaSeq v1.5, you also need to add an Index Read 2 custom primer oJR440. This is not necessary for MiSeq/NovaSeq v1 due to the grafted P5.
- PhiX Spike-in: Not necessary for libraries containing many guide RNAs with precise read lengths.

**Appendix 1. Detailed genomic DNA extraction protocol**

Materials per sample (up to 100 M cells)

- 1 column
- 500 uL proteinase K
- 10 mL BQ1
- 10 mL ethanol (+ extra to make BQ2)
- 15 mL BQ2
- 1.6 mL EB

Preparation:

- Preheat water baths to 56C and 70C
- Resuspend proteinase K in proteinase buffer PB (for NucleoSpin Blood XL: 5.75 mL per tube)
- add ethanol to buffer BQ2 concentrate
- heat buffer EB to 70C
- label 1 falcon tube, 1 column tube, 1 collection tube for each sample

Step-by-step protocol:

1. Thaw cells: RT (or 37C)
2. Transfer cells to 50 mL falcon tube with 25 mL PBS.
3. Pellet cells. Spin @ 1300 rpm (~400 x g), 5 min.
4. Resuspend each pellet in 10 mL PBS.
5. Add 500 uL proteinase K to each tube.
6. Add 10 mL BQ1 to each tube. Mix by vortexing vigorously (or by inverting 15 times).
7. Incubate at 56C for 15 min.
8. Cool to RT for at least 30 min. **THIS IS CRITICAL**
9. Add 10 mL 100% EtOH to each tube, mix by vortexing vigorously (or by inverting 15 times).
10. For each lysate, load 15 mL onto one column.
11. Spin 3 min at 4000 x g, discard flow-through. (or keep in tube)
12. Load remaining lysate onto column.
13. Spin 3 min at 4000 x g, discard flow-through.
14. Add 7.5 mL BQ2 to each column.
15. Spin 2 min at 4000 x g.
16. Add 7.5 mL BQ2 to each column.
17. Spin 10 min at 4000 x g.
18. Move column to a new collection tube.
19. Add 800 uL pre-heated buffer BE to each column.
20. Incubate at RT 5 min, then spin 4 min at 4000 x g.
21. Repeat steps 19+20.
22. Measure DNA concentrations (transfer 2 uL to a fresh tube, measure on NanoDrop).

**Appendix 2. PCR amplification of plasmid DNA rather than genomic DNA.**

We generally set up 5X 100 uL PCRs per library. We do not recommend exceeding an input of 10 ng pDNA per 100 uL reaction, as excess template DNA can lead to PCR-based uncoupling of guides (Hegde *et al*. 2018, PMID 29799876).

### PCR parameters

| Reagent | uL per 100 uL reaction |
| --- | --- |
| oJR441 reverse primer (100 uM) | 1 |
| Indexed forward primer (100 uM) | 1 |
| NEBNext Ultra II Q5 MasterMix | 50 |
| plasmid DNA | 10 ng |
| Water | bring to final volume of 100 uL |

### PCR conditions

- 98 ºC, 30 s
- 12-14 cycles (see note below)
  - 98 ºC, 10 s
  - 66 ºC, 75 s
- 72 ºC, 5 min
- 4 ºC hold

Note on number of cycles: empirically, 13 cycles produce sufficient amount of product. To be absolutely certain that enough PCR product will be obtained, cycle number can be increased to 14, with the caveat that each additional cycle may induce non-linearities or PCR uncoupling.

After the PCR is completed, pool all your reactions for a given sample. Mix reaction well. All of the sgRNAs in the pool should be represented at much greater abundance than, so you may proceed with only a fraction of the sample for PCR purification. Remember that this material represents a significant contaminant to pre-PCR samples, so be sure to keep them separate.
