## Supplementary material for "Maximizing CRISPRi efficacy and accessibility with dual-sgRNA libraries and optimal effectors": Supplementary_Note_3.docx

**Supplementary Note 3. Arrayed cloning of dual sgRNAs into lentiviral vectors**

This protocol works for low or medium throughput cloning of programmed pairs of sgRNAs into a lentiviral expression vector. The guide expression cassette of mU6-CR3-hU6-CR1 uses different U6 and sgRNA constant regions in the two positions in order to prevent intramolecular lentiviral recombination.

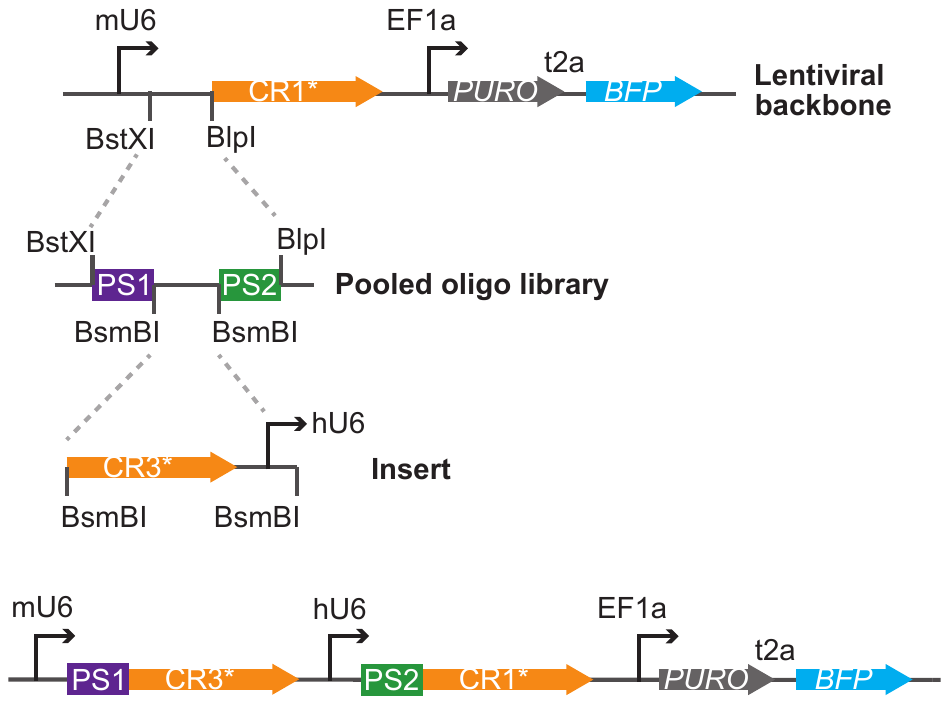

The strategy uses four components:

1) sgRNA expressing lentiviral vectors (note: it is not necessary to use a BsmBI- vector for this protocol. This feature is only important for pooled dual-guide library cloning.)

| **plasmid** | **U6 promoter** | **constant region** | **capture sequence** | **Pol-II promoter** | **Selectable marker** |
| --- | --- | --- | --- | --- | --- |
| pJR85 | mU6 | modified CR1 | CS1 in stem loop | EF1alpha  (no UCOE) | PURO-BFP (BsmBI-) |
| pJR100 | mU6 | modified CR1 | CS1 in stem loop | UCOE-EF1alpha | PURO-BFP  (BsmBI-) |
| pJR101 | mU6 | modified CR1 | CS1 in stem loop | UCOE-EF1alpha | PURO-GFP  (BsmBI-) |
| pJR103 | mU6 | CR1 | none | UCOE-EF1alpha | PURO-BFP  (BsmBI-) |
| pJR104 | mU6 | CR1 | none | UCOE-EF1alpha | PURO-GFP  (BsmBI-) |

2) top and bottom oligos annealed for position A sgRNA targeting sequence (see excel template)

3) CR3-hU6 insert

| **plasmid** | **U6 promoter** | **constant region** | **capture sequence** |
| --- | --- | --- | --- |
| pJR89 | hU6 | modified CR3 | CS1 in stem loop |
| pJR97 | hU6 | CR2 | none |
| pJR98 | hU6 | CR3 | none |

4) top and bottom oligos annealed for position B sgRNA targeting sequence (see excel template)

Capture sequences are used for 3’ direct capture Perturb-seq. For all other experiments, it is not necessary to use capture sequences in the sgRNA constant region. If you are cloning many sgRNAs, I recommend buying your oligos in pre-suspended at 100uM in a plate in an arrayed format that is easy to combine top and bottom oligo by multichannel. Currently, only ~30-50% of colonies contain the desired sequence (although I am working on troubleshooting this by either adjusting the ratio of vector to insert or using phosphatase treated vector).

### Step 1: Annealing of sgRNA targeting sequences:

48μL IDT duplex buffer

1μL	100uM top oligo 1μL	100uM bottom oligo 50μL total reaction volume

Anneal top and bottom oligo for position A. Anneal top and bottom oligo for position B. Incubate at 95°C for 5min in a PCR machine. Let oligos gradually anneal while cooling to RT (take the plate out and set it on your bench or use a cooling ramp on a PCR machine to cool the plate).

Make a 1:20 dilution of annealed oligos in ddH2O.

Annealed oligos can be stored a -20°C and are stable through at least 2-3 freeze thaws.

### Step 2: Insert digest

5ug	pJR98

10μL	10x 3.1 buffer

5μL	NEB BsmBI

xx μL ddH2O

100μL total reaction volume

### Incubate at 55 C for 1+ hour. Gel extract the desired 400 bp insert.

### Step 3: Vector digest

5ug	pJR103

10μL	10x 2.1 buffer

4μL	NEB BlpI

4μL	NEB BstXI

xx μL ddH2O

100μL total reaction volume

### Incubate at 37 C for 1+ hour. Gel extract the desired 9kb vector.

### Ligation:

100ng digested vector backbone (digested with BstXI and BlpI)

20ng digested insert (digested with BsmBI)

6μL	1:20 diluted annealed oligo pair A

6μL	1:20 diluted annealed oligo pair B

2μL	fresh 10X T4 ligase buffer 1μL	T4 ligase

xx μL ddH2O

20μL total reaction volume

Incubate at RT for 1-4hrs or 16C overnight.

Transform to DH5α bacteria:

Pre-warm LB agar + amp or carb plate(s) at 37°C. Combine 20μL cells + 1μL ligation; incubate 30min on ice.

Heat shock cells at 42°C for 45 sec (this can be done in tubes with a water bath or a PCR machine with 96 well plates- prewarm the PCR block to 42C and then set the plate in).

Put immediately back on ice for ~1 min.

For low throughput cloning

add 100μL LB to transformation and recover at 37°C shaking for 1hr. Plate 50μL of the transformation

Incubate at 37°C overnight.

For higher throughput cloning

Immediately plate all 20uL of transformed bacteria onto a spot on a partitioned plate and streak off the spot to single cell dilution. I use as little as 4uL of bacteria to save money.

Pick 4 colonies per sgRNA pair and miniprep (use Zymo Research Zyppy-96 (D4041) for 96 well plates minipreps format). Or use standard mini preps to prep plasmid DNA. As stated above, only ~30-50% of colonies contain the desired sequence although I am working on troubleshooting this by either adjusting the ratio of vector to insert or using phosphatase treated vector.

Sanger sequence with oBA420 (5’- cagcacaaaaggaaactcacc -3’)
