## Supplementary material for "Maximizing CRISPRi efficacy and accessibility with dual-sgRNA libraries and optimal effectors": Supplementary_Note_4.docx

**Supplementary Note 4: Generating CRISPRi cell lines**

Adapted and expanded from <https://weissman.wi.mit.edu/resources/CRISPRiacelllineprimer.pdf>

**Overview**

Here we describe our general strategy for generation of CRISPRi cell lines that mediate strong and consistent knockdown of target genes. The strategy consists of introducing an expression construct for a CRISPRi effector by lentiviral transduction, selecting a pure population of effector-expressing cells, and assaying knockdown. Importantly, these approaches are primarily optimized for dividing human cell lines, which are relatively stable in cell culture, rather than for organoids, primary cells, or stem cells, which require additional considerations.

The three key aspects in our experience are:

1) identifying a construct that allows for stable expression of a CRISPRi effector

2) selecting a very pure population of cells expressing the CRISPRi effector

3) Benchmarking knockdown using direct measurements of mRNA or protein levels

**1. Identifying CRISPRi effector constructs**

Our effector of choice is Zim3-dCas9, for the reasons outlined in the manuscript.

(Note that Zim3-dCas9 and other KRAB-containing effectors require endogenous TRIM28 to mediate knockdown. We have found Zim3-dCas9 to mediate knockdown across many cell types, but it may be worth testing a non-KRAB effector, such as MeCP2, in parallel if TRIM28 expression is undetectable in a cell type of interest.)

To select an expression construct, we consider the following criteria:

a. Promoter: We generally start with the SFFV and EF1a promoters, as one of these two confers strong expression in most cell lines. The SFFV promoter is small and very strong but gets silenced or is toxic in some cell lines. The EF1a promoter is bigger but mediates more consistent expression across cell types in our experience. In the absence of other information, we generally test both the SFFV and EF1a promoters. For cell types in which specific promoters are known to perform well, these promoters can be subcloned into our expression constructs. (In addition to the EF1a promoter, the EF1a-short/EFS promoter is also worth testing, as the EF1a promoter contains an intron that gets removed during lentivirus packaging.)
b. Ubiquitous chromatin opening element (UCOE): We have found that including a UCOE helps prevent silencing of effector expression and is essential for long-term stable expression in primary cells such as fibroblasts (e.g. PMID 34697476 and 27984733), consistent with previous reports (e.g. PMID 20588258 and 25605798,). Our default approach now is to include a UCOE upstream of the promoter.
c. Expression marker: We generally use fluorescent protein (FP) markers to enable selection for cells that express the effector by FACS. We have generated constructs that use BFP, GFP, or mCherry. Note that our pooled single sgRNA libraries (Horlbeck et al. 2016) are in BFP-marked vectors, which are more challenging to use together with a BFP-marked effector, while our pooled dual sgRNA libraries described here are in GFP-marked vectors. We generally use constructs in which the FPs are linked via a P2A ribosome skipping sequence, which give rise to high FP levels and thus make it easy to select cells by FACS. We also have constructs in which the FPs are linked via an IRES or constitutive fused to the effector, both of which lead to lower FP levels. Direct FP fusion may impact effector activity, although we have not identified cases in which this is an issue. For cases in which it is important to keep multiple fluorescence channels available, we have constructs in which the FP is flanked by LoxP sites and thus can be removed by transient delivery of Cre (pNM1130, pNM1129) or in which effector expression is marked with a hygromycin resistance gene (pAG389), which allows for selection of a pure population but may require repeated selection. In the remainder of this protocol we discuss cell line generation using FPs.

**2. Selecting a pure population of effector-expressing cells**

We introduce the expression construct for the CRISPRi effector by lentiviral transduction, followed by FACS with a stringent gate to select a pure population of effector-expressing cells. (Note: lentiviral constructs get silenced in some cell types; see the notes at the end for potential solutions.) We usually perform a second sort a few days after the first sort to ensure purity.

Brief protocol:

a. Generate lentivirus for the effector expression construct, using standard protocols. (e.g. transfect HEK293T cells at ~70% confluence with the expression construct and lentivirus plasmids, harvest culture supernatant with lentivirus 2-3 days after transfection.)

Notes:

- HEK293T health is critical to obtaining high-titer lentivirus. Use cultures of HEK293T cells that have been maintained at below 70% confluence for several passages.
- It is a good idea to package lentivirus at the scale of a 10-cm or a 15-cm plate.
- The expression constructs are big, such that lentiviral titers may be low. If needed, concentrate the virus by precipitation or ultracentrifugation (usually not necessary).
- To minimize carryover of HEK293T cells, consider freezing the lentivirus or filtering the lentivirus through a 0.45 µm syringe filter during harvesting. The lentivirus can be aliquoted and stored at −80 C for weeks to months. Assume that titers will drop ~50% upon freeze-thawing.

b. Transduce ~1-3 x 10^6^ target cells with lentivirus at a MOI of 0.2-1 (~20-50% cells infected).

Notes:

- For most cell types, transduction with 100-2000 µL virus per 1 x 10^6^ cells in the presence of 8 µg/mL polybrene or a similar cationic polymer will yield the expected MOI. Spin-infection at 1000 x g and 33 C for 0.5-2 h can help boost transduction.
- When working with a new cell type and/or a new batch of lentivirus, titrate the lentivirus by infecting cells with different volumes of lentivirus and measuring transduction efficiencies by flow cytometry.

c. Select effector-expressing cells by FACS 3-5 days after transduction. Sort 2 x 10^5^ – 2 x 10^6^ transduced cells using stringent FP and singlet gates. After the sort, set aside 1-5% of the sorted cells and immediately run them on a flow cytometer for a purity check. You want essentially 100% of cells to be FP-positive (>99.9%).

Notes:

- Ensure that the cells for FACS are well-dissociated to minimize doublets. Strain the cells to remove clumps/aggregates.
- Use an untransduced control population to set a stringent area gate (e.g. mCherry vs. BFP or SSC) that excludes 100% of the untransduced population. Consider moving the gate several AUs away from the untransduced population to minimize background.
- Set the sorter to high-purity mode, if available.
- Plan to sort at least 2 x 10^5^ cells and ideally more. Sorting many cells minimizes the impact of any single lentivirus integration site in the polyclonal population.

d. Recover and expand the sorted cells using standard protocols. For some cell types, it can help to seed the cells at elevated density, with 20% FBS, and/or with 20% conditioned media.
e. **Our standard approach is to perform a second sort**, especially if post-sort purity was <99%. In this case it is sufficient to expand the cells for 3-5 days. Then re-sort using the same protocol. (This sort will be a lot faster as the input population should be >90% FP-positive, so sorting 2 – 5 x 10^6^ cells is feasible.)
f. Recover and expand the sorted cells. At ~10 x 10^6^ cells, set aside a small number of cells to assess knockdown. Continue expanding the remaining cells to 50-200 x 10^6^ cells and freeze aliquots using standard protocols. Aim to freeze >20 aliquots so that you can return to the originally sorted population whenever needed.

**3. Benchmark knockdown**

We have found it important to directly measure knockdown of target genes, as opposed to measuring phenotypes resulting from knockdown, such as growth defects of essential gene knockdowns. The main reason is that we generally consider >95% knockdown to be good, but for many essential genes, 50-80% knockdown is sufficient to confer a growth phenotype. Thus, the ability to measure a growth phenotype does not guarantee good knockdown.

The simplest approach to measuring knockdown is to express sgRNAs against non-essential cell surface proteins and then directly measure protein levels by flow cytometry. The benefits of this approach are 1) a straightforward protocol, 2) no need to select for sgRNA-expressing cells as you can simply gate for sgRNA-expressing cells, and 3) the ability to detect subpopulations of cells without knockdown in the single-cell data. This approach consists of the following steps:

a. Identify cell surface markers that are expressed in the cell type of interest. We generally work with some combination of CD55, CD81, CD151, and B2M, all of which are non-essential, near-ubiquitously expressed, and have good antibodies available (Methods).
b. Select sgRNAs targeting these genes and a non-targeting control sgRNA, for example from Table S7 in this manuscript. Clone expression vectors for these sgRNAs. Use a backbone with a compatible fluorescent marker (e.g. BFP if your effector is marked with mCherry).
c. Generate lentivirus for the sgRNA expression vectors.
d. Transduce ~1-2 x 10^5^ CRISPRi cells with the sgRNA expression vectors at a MOI of 0.2-1 and grow the transduced cells for 4-8 divisions (no selection necessary).
e. Stain the transduced cells with fluorescently-labeled antibodies using standard protocols and use flow cytometry to measure fluorescence levels.

Notes:

- Selection for sgRNA-expressing cells is unnecessary. Simply gate for sgRNA-expressing cells in the flow cytometry data using the FP marker.
- Growing the cells for 4-8 divisions post-transduction ensures that protein levels reach their new steady state. As the targeted proteins are non-essential, there should be minimal selection against cells with knockdown.
- For new cell types, titrate antibodies to ensure detection in the linear range. A good starting point is to titrate the antibody across a range of 0.5 to 5 µL antibody per 100 µL cells. (The key parameter is antibody volume per cell volume.)
- To improve accuracy, consider also staining cells using an isotype control antibody, at least for cells expressing a negative control sgRNA. Subtract the signal from this antibody to more accurately quantify target protein levels.
- Use a fluorescent conjugate that is compatible with the FPs in your experiment. APC works well in most cases.

As an alternative approach, we also use qPCR to measure mRNA levels of non-essential genes after knockdown. The workflow is similar to the one described above, except that we select the sgRNA-expressing cells using FACS or an antibiotic, as qPCR is a bulk readout. Our default sgRNA expression vectors contain a puromycin resistance gene. We generally start selection 2-3 days after transduction, and then select for 3-5 days until >99% of cells are sgRNA-expressing, as assessed by flow cytometry. We let the cells recover for 1-2 days, harvest the cells, extract RNA, and perform qPCR using standard protocols. The table at the end contains sequences of potential sgRNAs.

**Additional Notes**

1. Constitutive versus inducible expression

In this protocol we only discuss constitutive expression of the CRISPRi effector. We and others have generated constructs for inducible expression, e.g. using the Tet system (Gilbert et al. 2014, Mandegar et al. 2016, Tian et al. 2019, Tian et al. 2021), anti-CRISPRs (Nakamura et al. 2019), or degrons (Tian et al. 2019, Tian et al. 2021). In most cases, especially in cancer cell lines, constitutive expression covers most applications and is simpler to implement because it only requires a single component. That said, in some cases an inducible CRISPRi system may be beneficial, which can be generated using the general approach outlined here.

2. Selecting for high-knockdown cells using an activity-based sort

In some rare cases, subpopulations of cells do not have knockdown despite expressing the effector. In these cases it can be beneficial to perform an activity-based sort that directly selects for cells with strong knockdown:

a. Transduce cells with the effector expression construct as described above.
b. Transiently deliver an sgRNA against a non-essential cell surface protein (e.g. the ones described above), for example by electroporation or transfection of the purified sgRNA.
c. Allow the cells to grow for ~3-5 days.
d. Stain the cells with a fluorescently-labeled antibody against the targeted protein and then perform FACS with an additional gate for cells that have knockdown of the targeted protein (i.e. low antibody staining intensity).
e. Perform remainder of cell line generation as described above.

- Note: It is important that the sgRNA is only present transiently, so that the sorted cells do not permanently express the sgRNA. The simplest approach is to electroporate or transfect the purified sgRNA (e.g. commercially obtained). An alternative is to use non-integrating AAV vectors. In some cases it may be necessary to transiently transfect an sgRNA expression plasmid. In this case it is critical to ensure that the sorted cell population does not contain cells that integrated the plasmid, for example by genomic qPCR.

3. Polyclonal vs. monoclonal populations

Our protocol is designed to select for polyclonal populations of cells. As described above, we ensure that knockdown is homogeneous across the cell population by measuring knockdown at the single-cell level. In case there are subpopulations of cells without knockdown, we first use the activity-based sort. Only if we are unable to eliminate subpopulations of cells without knockdown do we generate clonal CRISPRi cell lines, with the same approach but sorting for single cells. The main caveat of clonal cell lines is that the clones can differ in many aspects from the parental population, including drug sensitivity, mutation status, and others, such that it is critical to validate the biology of interest in the clone and to ensure that any genetic hits are not due to idiosyncrasies of the clone. In addition, we have observed that even clones selected from a polyclonal population with homogeneous knockdown can have variable knockdown, so it is equally important to validate that clones still retain stong knockdown.

4. Assessing and counteracting silencing

In some cell types, the lentiviral expression construct gets silenced over time, which leads to loss of CRISPRi activity. This is one of the reasons to freeze as many aliquots as possible shortly after FACS. When using cells from later passages, it is a good idea to re-test for CRISPRi activity using the approaches outlined above. (Note that low levels of the fluorescence protein marker do not necessarily indicate lack of CRISPRi activity, which is why we recommend directly assaying CRISPRi activity.) In cell types with immediate and severe silencing, we recommend considering 1) piggyBAC transposon-based systems (note that piggyBAC systems often integrate multiple copies of the payload, which may result in higher expression of the CRISPRi effector; we have not tested comprehensively if such high expression results in non-specific effects on cell growth and transcription), 2) site-specific recombination-based integration into safe-harbor loci, such as AAVS1 (Mandegar et al. 2016); 3) periodic re-sorting of cell lines; 4) generating clones (see above); or 5) a combination of these approaches.

**Table of sgRNAs for quantification of knockdown at the mRNA level**

| Gene name | sgRNA name | sgRNA sequence | mRNA remaining (K562) | qPCR fw | qPCR rv |
| --- | --- | --- | --- | --- | --- |
| ST3GAL4 | ST3GAL4__ST3GAL4_w_126225631.24 | GGCGCGACGGCTCGACTCGGC | 0.030 | GATCACGCTCAAGTCCATGG | CTTGCCCAGGTCAGAAGGA |
|  | ST3GAL4__ST3GAL4_w_126225595.25 | GGGCGCGGGTCCGGCCTGGGAG | 0.019 |  |  |
| SEL1L | Apoptosis+Cancer+Other_Cancer=SEL1L_+_82000162.24-all~e39m1 | GCAGGAAGAGCAGCGGCGAGG | 0.008 | GAGGGGGAAAGTGTCACAGA | GGTCAAAGCTGGTTTCCGTA |
|  | Apoptosis+Cancer+Other_Cancer=SEL1L_-_82000023.22-all~e39m1 | GGGGGCGGATACTGACCCG | 0.024 |  |  |
| SYVN1 | Stress_Proteostasis=SYVN1_+_64901917.24-all~e39m1 | GGCGCTGGGTTCCTGGTGAGT | 0.022 | ACCAGCATCCCTAGCTCAGA | TCCTCAGGCATCTCCTCTGT |
|  | Stress_Proteostasis=SYVN1_-_64901760.22-all~e39m1 | GTTGCGGGCGTCGCAGGCA | 0.026 |  |  |
| UBE4A | Stress_Proteostasis=UBE4A_-_118230398.25-all~e39m1 | GCACTTCAGGCTCTGGTAAGAC | 0.024 | CAGCTGGAAAGCCTCAAATC | GGCATTTGCAAAAGTGTCCT |
|  | Stress_Proteostasis=UBE4A_+_118230531.23-all~e39m1 | GACAGGGTCGAAGTTCAAGA | 0.087 |  |  |
| DPH1 | Apoptosis+Cancer+Other_Cancer=DPH1_-_1933485.21-ENST00000263083.6~e39m1 | GGCGGCGCTGGTCGTATC | 0.026 | TGAAAGCCGAGTATCGTGTG | AGATACACAACGGCCTCCAC |
|  | Apoptosis+Cancer+Other_Cancer=DPH1_-_1933500.23-ENST00000263083.6~e39m1 | GTATCCGGGGCAGCGGAGCA | 0.017 |  |  |
| DPH2 | Gene_Expression=DPH2_+_44435896.24-all~e39m1 | GAGTAAGCAGTCCTGGCACCC | 0.033 | ACCTGGACGGAGTGTACGAG | TCTCCCAATAGCTGGTCAGG |
|  | Gene_Expression=DPH2_-_44435877.23-all~e39m1 | GATGTTTAGCAGCCCTGCCG | 0.095 |  |  |
| ST3GAL5 | Membrane_Proteins=ST3GAL5_-_86116115.23-all~e39m1 | gcTCTCGCGCCCATTCAGCT | 0.001 | GAAGGAGAGGGGAGGTCACT | AGAGCTACGGAGCACGTCAT |
| FUT4 | Membrane_Proteins=FUT4_-_94277315.24-all~e39m1 | GTAGCGGATGAGGCGCTTGTG | 0.016 |  |  |
|  | Membrane_Proteins=FUT4_+_94277081.24-all~e39m1 | GGACCCGGCGCCGCGGAACGG | 0.011 | CTTGGCTCCTGAGAAAGGTG | CAGCACTTTGCTTCCATTCA |
| H19 | H19_+_2019023.23-all~e39m1 | GGGGGGTAACGGGGGAAACT | 0.015 |  |  |
|  | H19_+_2019063.25-all~e39m1 | GCTAGGACCGAGGAGCAGGGTG | 0.013 | CTGGCTTGGCAGACAGTACA | TCCCTCCTGAGAGCTCATTC |
| XIST | XIST_+_73072490.25-all~e39m1 | GACTGAAGATCTCTCTGCACTT | 0.022 |  |  |
|  | XIST_+_73072528.24-all~e39m1 | GCCATATTTCTTACTCTCTCG | 0.016 | AGGTCAGGCAGAGGAAGTCA | CTGCCTCCCGATACAACAAT |
